# *Leishmania*-infected human dendritic cells clear their intracellular parasites by a mechanism involving the cytolytic effector molecules granzyme B and granulysin

**DOI:** 10.64898/2026.10.08.757588

**Authors:** Ibrahim Mohamed Azzouz, Renato Brito Baleeiro, Lydon Wainaina Nyambura, Arkadi Kundik, Peter Walden

**Affiliations:** Charité – Universitätsmedizin Berlin, corporate member of Freie Universität Berlin, Humboldt-Universität zu Berlin, and Berlin Institute of Health, 10098 Berlin, Germany; London School of Hygiene & Tropical Medicine (LSHTM), Faculty of Infection and Tropical Disease, Department of Infection Biology (Honorary Professor), Keppel Street, London WC1E 7HT, United Kingdom

**Author notes:** Corresponding author: Peter Walden. These authors contributed equally to the work (Ibrahim Mohamed Azzouz, Renato Brito Baleeiro, Lydon Wainaina Nyambura).

**Keywords:** adaptive immunity, dendritic cell, macrophage, granzyme, granulysin, infectious disease, innate immunity, Leishmania, phagocyte, T cell, vaccine

## Abstract

*Leishmania* are intracellular parasites that cause various human diseases ranging from self-healing cutaneous to fatal visceral leishmaniasis. Host cells are phagocytes, primarily macrophages, where the parasites neutralize innate immune defenses, proliferate and finally rupture the cells. Despite that, *Leishmania* infection induces vigorous T cell responses, which require antigen processing and presentation, and stimulation by phagocytes, importantly dendritic cells. It was not clear how to align blockade of phagocyte functions with efficient immune stimulation. Here we show that, in contrast to macrophages, human dendritic cells survive the infection and destroy the parasites. This process involves granzyme B and granulysin, two cytolytic molecules essential for destruction of infected or otherwise aberrant cells by cytolytic T cells and natural killer cells. The report for the first time shows that these molecules not only act in target cell lysis by cytolytic effector cells but also in intracellular processing of infectious agents by dendritic cells.

## Introduction

Leishmaniases are vector-borne infectious diseases caused by protozoan parasites of the genus *Leishmania*, which are transmitted by female phlebotomine sandflies during blood meal^1^. Depending on parasite species, vector and immune state of the infected individual, the clinical manifestations vary greatly. The three major forms are cutaneous, mucocutaneous and visceral leishmaniasis. The clinical manifestations of cutaneous leishmaniasis include skin nodules, ulcers or blackened maculae. Mucocutaneous leishmaniasis is characterized by ulcers of mucosa, especially in mouth and nose. Visceral leishmaniasis, the most severe form, causes splenomegaly, hepatomegaly, wasting and bouts of fever, and is fatal if not treated^2–4^. Leishmaniases are most frequent in the tropics, subtropics and some areas of southern Europe^3,5^.

Sandflies are pool feeders that disrupt blood vessels and suck from the resulting blood pool. Thereby the promastigote *Leishmania* parasites are immediately exposed to white blood cells and rapidly engulfed by neutrophils and macrophages. Once inside the host cells, the parasites convert to the amastigote state and undergo multiple cell divisions, eventually disrupting the phagocyte membrane and infecting new macrophages^6^. The parasites have developed effective strategies to evade the macrophage effector functions such as adapting to the acidic conditions in the phagolysosomes and altering their glycocalyx coat to protect their membrane in order to survive and multiply^6^. *Leishmania* parasites can be destroyed inside infected macrophages by cytolytic T cells by granulysin-mediated up-take of granzyme B that cleaves, among others, respiratory enzymes resulting in accumulation of active oxygen species and death of the parasites^7^. This process appears to be the major mode of parasite clearance in human macrophages but not in mice as they lack granulysin^8^. Although macrophages are regarded as most important host cells, fibroblasts, neutrophils, and dendritic cells (DCs)^9^ have also been shown to harbor *Leishmania* parasites. However, to date little is known about the impact of *Leishmania* infection on DC functions.

DCs are highly specialized cells that functionally link innate and adaptive immunity^10^. They are major antigen presenting cells (APCs) and distributed in all peripheral tissues of the body in an immature state, where they are highly efficient in taking up particulate and soluble material, and processing it for presentation by their MHC class I and class II molecules. When triggered by microbial factors, inflammatory cytokines or cognate interaction with T cells, DCs undergo a maturation process characterized by up-regulation of MHC class I and II molecules, the co-stimulatory molecules CD40, CD80 and CD86, the adhesion molecule CD58, *de novo* expression of CD83 and production of IL-12. These changes enable DCs to efficiently induce T cells and thereby adaptive immunity^11^.

Despite the well-established capacities of DCs as phagocytes, link between innate and adaptive immune systems and inducers of cellular immune responses, the effect of *Leishmania* infection on the functions of human DCs is largely unexplored. The infection induces vigorous *Leishmania* antigen-specific CD4+ and CD8+ T cell responses, which suggests a prominent role of DCs in the processes^12–14^. A better understanding of how *Leishmania* affect DC functions could help to improve DC responses for the control of *Leishmania* infection and development of vaccines. We investigated the fate of *L. donovani* inside DCs, and the effect on human DC maturation and induction of CD4+ and CD8+ T cells.

## Results

### Uptake and processing of *L. donovani* by human macrophages and DCs

We monitored uptake and intracellular routing of *Leishmania* parasites by human DCs with YFP-transfected live or killed *L. donovani*. Day 5 immature DCs (iDC) and for comparison day 8 macrophages differentiated from human peripheral blood monocytes by culturing in the presence of GM-CSF plus IL-4 or GM-CSF alone, respectively, were infected with promastigotes at a MOI of 10:1 or equivalent number of dead parasites. The uptake of the parasites by macrophages was more efficient with live than with dead parasites. At 2 h the percentage of cells with live parasites was 19% compared to 11% with dead parasites. The percentage with dead parasites steadily declined with time to 1.9% at 96 h. The percentage with live parasites increased to 22% at 24 h and remained high until 96 h (Fig. 1A and B). DCs were more efficient in taking up parasites, especially dead ones. Already after 30 minutes, about 60% of the DCs had taken up dead and about 30% live parasites. The frequency of DCs with dead parasites increased up to 6 h and then declined. The fraction with live parasites still increased up to 24 h before decreasing to 12% after 96 h (Fig. 1C and D).

**Figure 1.**
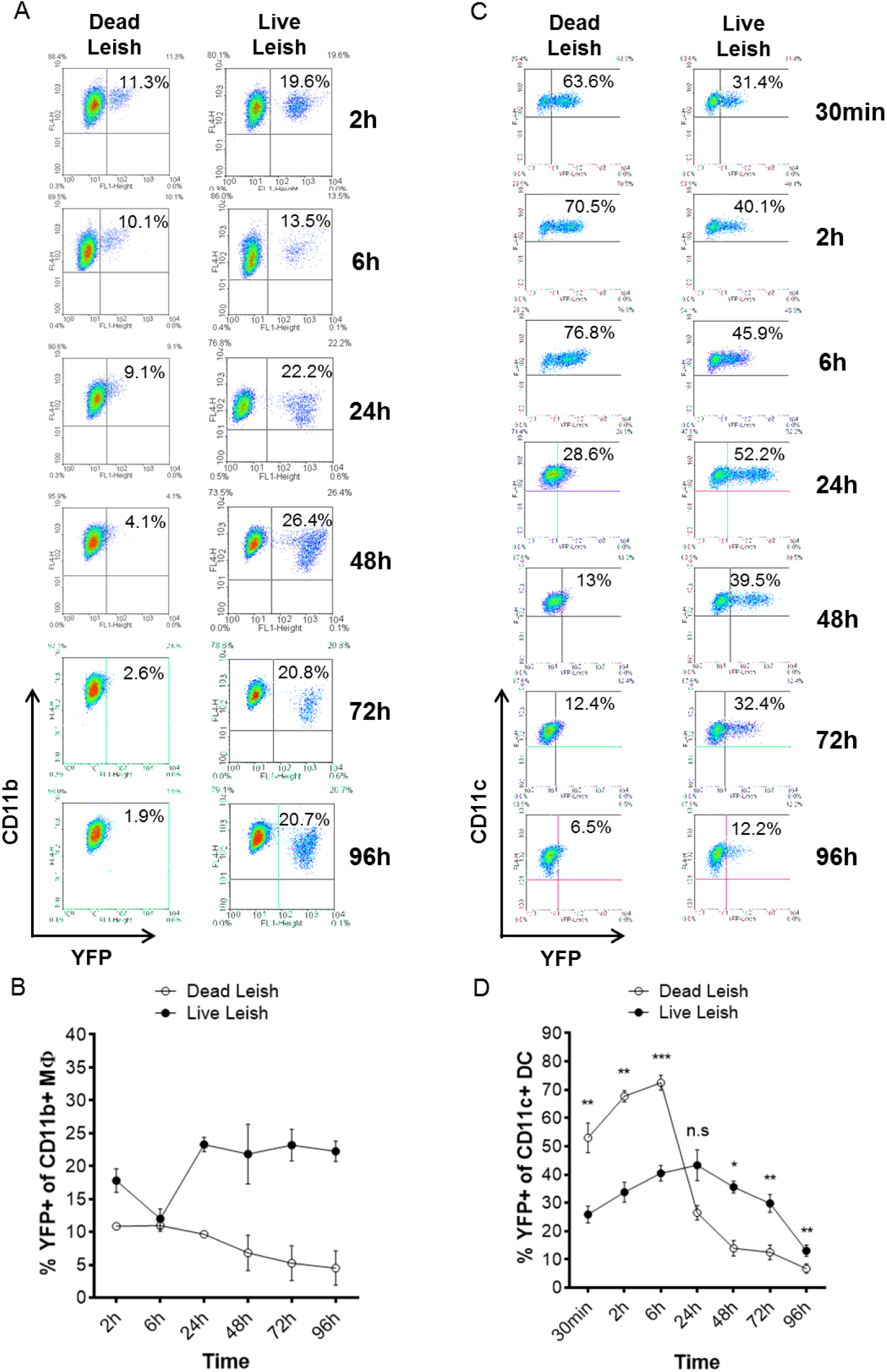
Kinetics of uptake of *L. donovani* parasites by human macrophages and iDCs. Day 8 macrophages and day 5 iDCs were incubated with dead or live YFP-*L. donovani* parasites, harvested at the indicated time points, stained with fluorochrome-conjugated anti-CD11b or anti-CD11c antibodies, respectively, and analyzed by flow cytometry. Data displayed in (**A**) are representative of two independent experiments performed with MΦ from two different donors. The graph in (**B**) shows mean of the percentage of CD11b+ that took up Leishmania parasites; n= 2. Data displayed in (**C**) are representative of three independent experiments performed with iDCs from three different donors. The graph in (**D**) shows mean of the percentage of CD11c+ that took up *Leishmania* parasites; n= 3. Error bars correspond to the standard error of the mean. n.s= non-significant; * = p<0.05; ** = p<0.01; and *** = p<0.001 comparing the group “Dead Leish” vs “Live Leish” at the indicated time points.

To identify the subcellular parasite-harboring compartments in DCs and test their relevance for antigen presentation and T cell stimulation we stained DCs incubated with live or dead parasites with fluorochrome-labelled antibodies against HLA class I and class II, the antigen-presenting molecules for CD8+ and CD4+ T cells, respectively, the invariant chain CD74, which is associated with newly biosynthesized but not recycled HLA class II to stabilize the molecule and protect the peptide binding groove, and calnexin, a membrane protein in the endoplasmic reticulum (ER) that acts as chaperon for nascent HLA class I. The invariant chain is regularly translocated together with the HLA class II into the phagosomes. Calnexin mostly remains in the ER but may also translocate into phagosomes for Ag cross-presentation, an important process in the induction of cytotoxic T cells^15,16^. Both live and dead parasites were routed into compartments positive for HLA class I and II, invariant chain CD74 and calnexin (Fig. 2). Already 2 h after addition to the cultures, the dead parasites were found in such compartments. For live parasites, the translocation appears to be slower. Thus, *Leishmania*-containing phagosomes expressed major components of the HLA class I as well as class II epitope-loading machinery. Moreover, the compartment containing the parasites become acidified as determined with LysoTracker, and over time the parasites are fragmented (Fig. S2). Taken together, these results suggest that, while Leishmania parasites survive in macrophages, in DCs they are routed into compartments associated to antigen processing for presentation by HLA class I and II and are destroyed.

**Figure 2.**
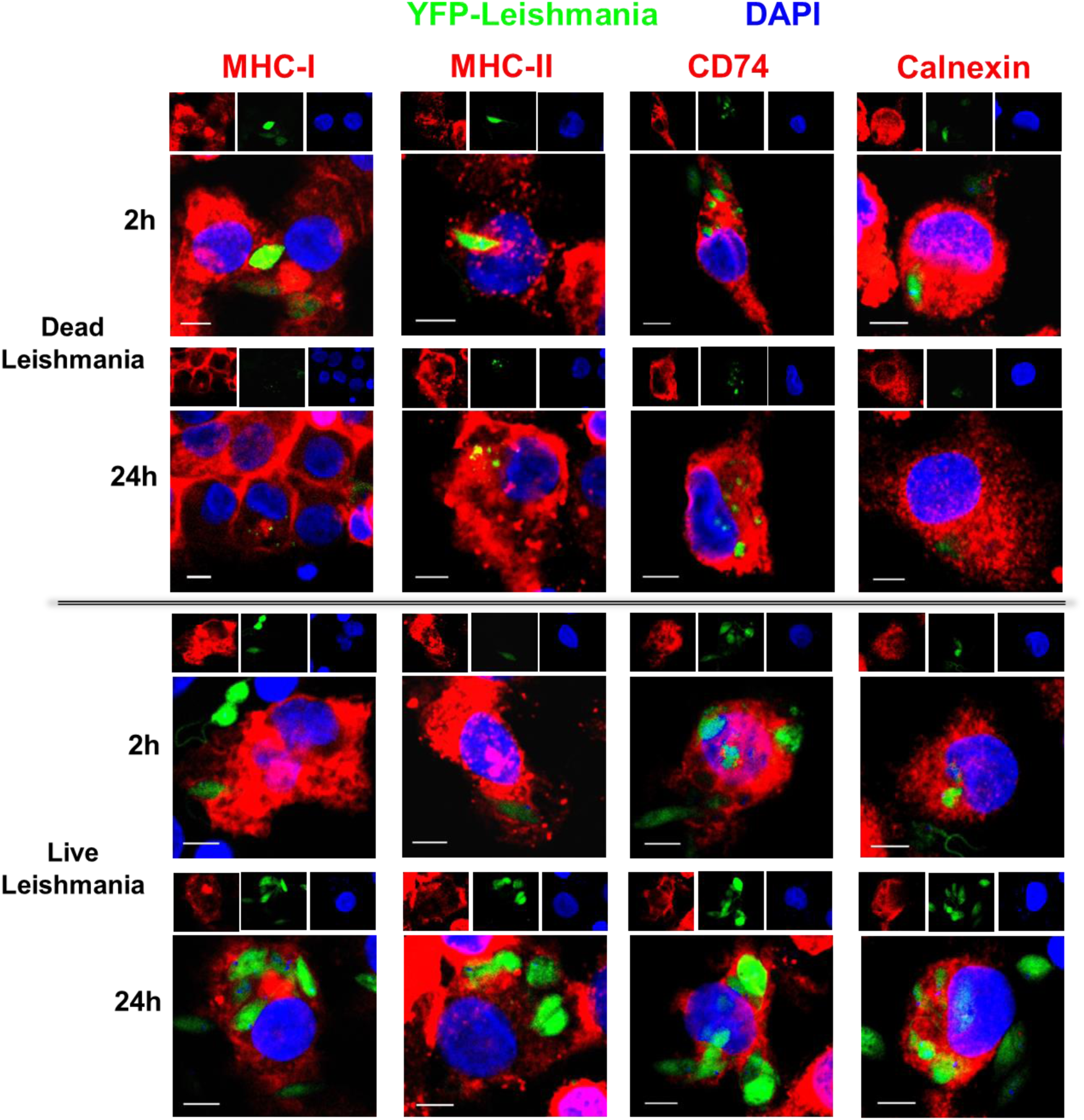
Intracellular fate of *L. donovani* parasites following uptake by human iDCs. Day 5 iDCs were incubated with dead or live YFP-*L. donovani* parasites (green fluorescence), harvested at the indicated time points, fixed, permeabilized and stained for MHC class I, MHC class II, CD74 and calnexin (red fluorescence), and counterstained with DAPI (blue fluorescence). After staining, the cells were imaged by confocal microscopy. Yellow indicates the overlay of the green YFP fluorescence and the red fluorescence of the indicated marker proteins. Scale bar= 5 μm.

### *Leishmania* parasites do not affect DC viability

To control that the decrease of infected DCs is not due to cytolytic effects of the parasites and relative increase of DCs that are resistant to infection, we incubated DCs with parasites, harvested them at different time points and analyzed their viability with calcein-AM, a live cell stain, and propidium iodide (PI), a dead cell stain. Staining for CD11c was included to identify DCs. The viability of DC without parasites over 96 h decreases from about 98% to 93% (Fig. S1 and Fig. 3). About the same values were obtained for DCs incubated with dead and those infected with live *Leishmania* parasites. This indicates that the reduction of DCs bearing parasites (Fig. 1B) is not due to selection of DCs that are resistant to infection. This contrasts with the situation with other phagocytes like macrophages and neutrophils^9^ where the parasites proliferate and destroy their host cells.

**Figure 3.**
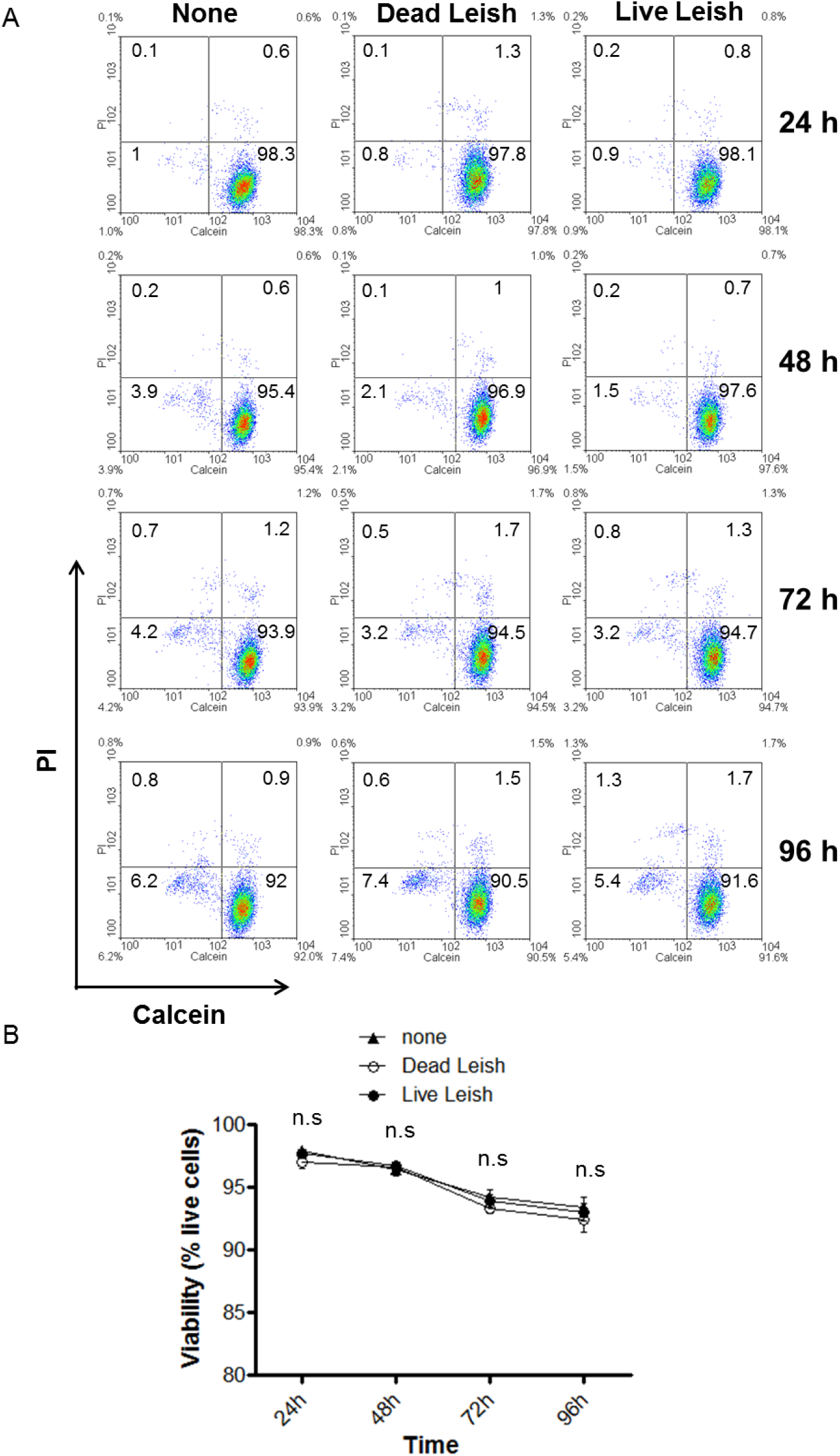
Effect of *L. donovani* on the viability of human iDCs. The viability of the DCs was assessed upon incubation with the parasites for the indicated times. DCs were then stained with the nonfluorescent cell-permeable calcein-AM which is metabolized in the cytoplasm of live cells into the green fluorescent calcein, and propidium iodide (PI), which only enters cells with perforated cell membranes and binds nucleic acids, staining the cells red. After staining, the DCs were analyzed by flow cytometry. Data shown in Figure S1 are representative of four independent experiments performed with cells from four different donors. The graph in Figure 3 shows mean of the percentage of calcein-positive cells. Error bars correspond to the standard error of the mean; n= 4. n.s= non-significant for the difference between the DCs without parasites versus those with dead parasites or infected with live parasites.

### Granzyme B/granulysin-dependent clearance of parasite in human DCs

Given the effective cytolytic capacity of *Leishmania*-infected human DCs against their intracellular parasites that macrophages lack, we compared published and archived transcriptome data for these two cell types via BioGPS. The analysis revealed high granzyme B expression by DCs compared to all other white blood cells, even higher than in cytolytic T cells, with low expression or virtual absence in macrophages (Fig. S3). Granzyme B is an essential cytolytic effector molecules of cytolytic CD8 T cells and natural killer cells ^17, 18^. Likewise DCs express granulysin, a saponin family molecule that is important for the destruction of intracellular bacteria and parasites by cytotoxic and natural killer cells ^19–25^ that macrophages also lack. We confirmed the differential expression of granzyme B and granulysin in macrophages and human DCs by RT-PCR (Fig 4A and F,and Fig. S4), and at the protein level by flow cytometry (Fig. 4B and G) as well as by confocal microscopy (Fig. S5). Granzyme B expression increased in DCs upon infection with *Leishmania* parasites (Fig. 4B and Fig. S5). Staining DCs infected with YFP-transfected parasites for 48 h with Alex Flour 647-conjugated anti-human granzyme B antibodies showed co-localized of fragmented parasites with granzyme B (Fig. 4C).

**Figure 4.**
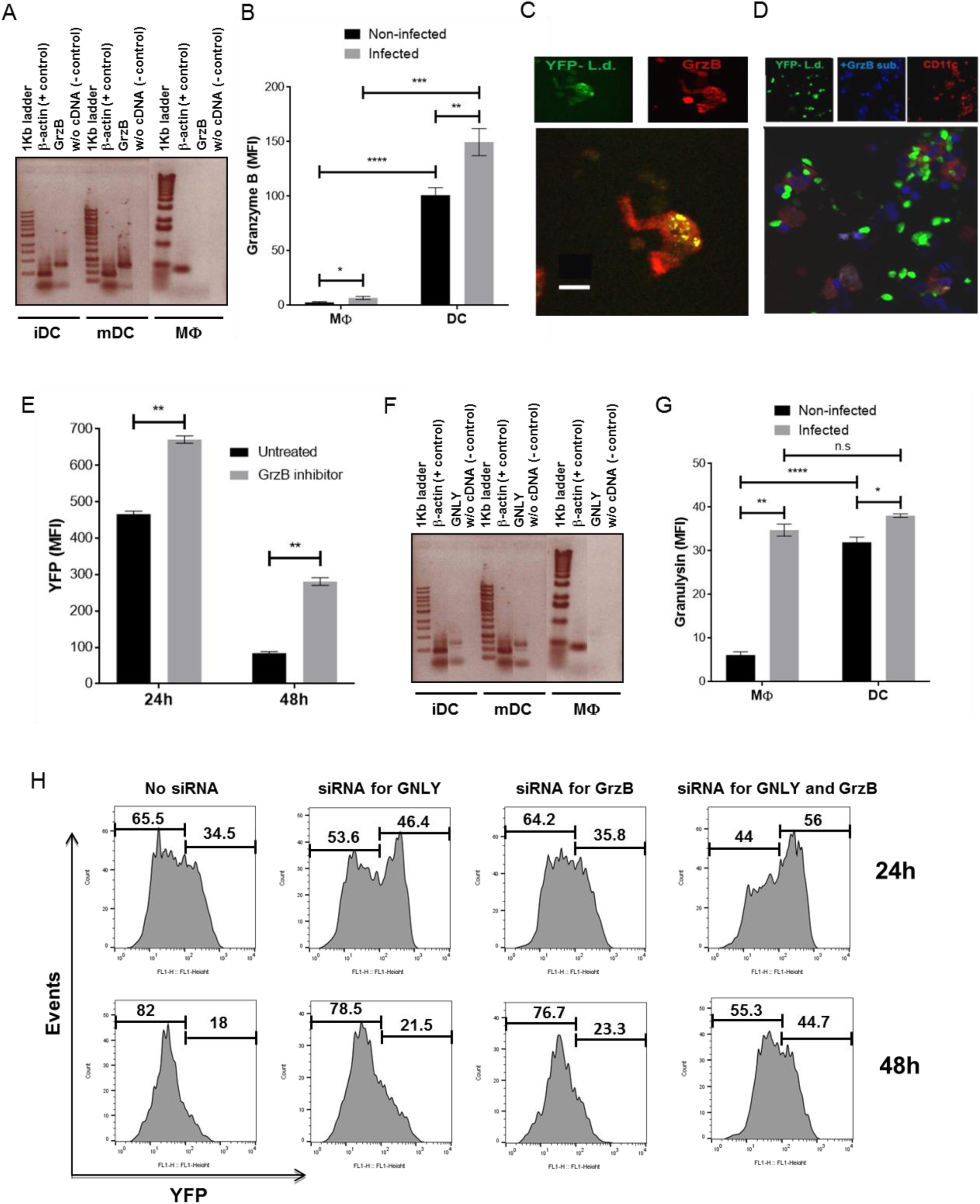
Granzyme B and granulysin in the elimination of *Leishmania* parasites by human iDCs. Day 8 macrophages and day 5 iDCs were incubated for 48h in absence or presence of live *L. donovani* parasites, harvested and processed for RT-PCR, flow cytometry and confocal microscopy as described in Material and Methods. (**A**) Results of RT-PCR for GrzBs in iDCs, mDCs and macrophages. β actin served as control for the PCR reactions, PCR with cDNA from PHA-activated PBMCs and PCR without cDNA as a positive and negative control, respectively. The black lines on the gels delineate grouping of gels cropped from different parts of the same gel, or from different gels (Full gels in Supplementary Fig. S4). (**B**) MFI of granzyme B protein expression in CD11b+ macrophages and CD11c+ DCs determined by flow cytometry; n=4. (**C**) Confocal image showing co-localization of *L. donovani* parasites (green YFP-fluorescence) with granzyme B (red fluorescence) in DCs. Yellow indicates an overlay of green and red fluorescence. Scale bar= 5 μm. (**D**) Confocal image indicating granzyme B activity in iDCs co-cultured with YFP-transfected parasites (green fluorescence) in the presence of fluorogenic granzyme B substrate for 48 h. Processing of the substrate by granzyme B generated a product with blue fluorescence. Red fluorescence marks CD11c on the cell surface of DC. (**E**) Effect of granzyme B inhibition on the clearance of *L. donovani* by DCs. Day 5 human iDCs were incubated with live YFP-*L. donovani* parasites in the absence or presence of granzyme inhibitor, harvested at the indicated time points, stained with fluorochrome-conjugated anti-CD11c and analyzed by flow cytometry. The mean fluorescence intensities (MFI) of YFP in iDCs gated on the CD11c+ population; n= 3. (**F**) Results of RT-PCR for granulysin mRNA in iDCs, mDCs and macrophages together with controls as in (**A**). (**G**) Granulysin protein expression in macrophages and DCs determined by flow cytometry as for granzyme B shown in (**B**); n= 4. (**H**) Involvement of granulysin and granzyme B in the clearance of *L. donovani* by DCs. Day 5 human iDCs were incubated with siRNA for GNLY, GrzB, GrzB and GNLY or left untreated and then infected with live YFP-transfected *L. donovani* parasites. The cells were harvested at the indicated time points, stained with fluorochrome-conjugated anti-CD11c and analyzed by flow cytometry. The histograms are representative of three independent experiments performed with cells from three different donors. Graphs in (**B**), (**E**) and (**G**) show mean values and the standard error of the mean; * p<0.05; ** p<0.01; *** p<0.001; and ****p< 0.0001.

The involvement of granzyme B in the clearance of intracellular *Leishmania* parasites by infected human DCs was substantiated by demonstrating granzyme B enzymatic activity in parasite bearing intracellular compartments by co-incubation with a granzyme B fluorogenic substrate for 48 h and confocal microscopy (Fig. 4D). Furthermore, incubation of DCs with parasites in presence of a granzyme B inhibitor resulted in enhanced survival of the intracellular parasites as demonstrated by enhanced YFP-positive DCs by flow cytometry (Fig. 4E). The treatment with the granzyme B inhibitor did not affect DC viability (Fig. S6).

The essential roles of granzyme B and granulysin in intracellular processing of *Leishmania* parasites by human DCs was confirmed by siRNA blocking experiments with YFP-transfected *Leishmania* parasites separately for granzyme B and granulysin and the combination of both (Fig. 4H). The cultures were analyzed by flow cytometry after 24 h and 48 h of infection in the presence or not of siRNA for YFP fluorescence in CD11c-positive cells. The siRNA treatments resulted in an enhancement of the frequencies of YFP-positive DCs whereby blockade of granulysin had a stronger effect than that of granzyme B, which was further enhanced by the combination. These observations are in agreement with the previously reported function of granulysin to mediate entry of granzyme B into the parasites for inducing parasite cell death through proteolysis of various essential proteins inside the parasites ^22^.

### *Leishmania* infection sensitizes DCs for enhanced inflammatory responses

DCs are essential for inducing strong and sustained T-cellular immune responses, and are activated by various microbial factor ^26, 27^. To examine the effect of *Leishmania* infection on the activation and maturation state of human DCs, DCs were differentiated from monocytes in the presence of IL-4 and GM-CSF for 5 days and then stimulated with LPS or a cocktail of the inflammatory cytokines IL-1β, IL-6 and TNF-α for 48 h in the presence or not of dead or live *Leishmania* parasites added two hours prior to LPS or the cytokine cocktail, so the DC could internalize the parasites prior to induction of maturation. Maturation was monitored by up-regulation of CD40, CD58, CD80, CD83, CD86, HLA-ABC, HLA-DR, down-regulation of CD1a and production of the biologically active IL-12, IL-12p70 ^11^. Incubation of the DCs with dead of live parasites did not alter the expression of these molecules (Fig. S7A-C). However, the addition of parasites followed by the proinflammatory cytokine cocktail enhanced DC activation, as evident from increased expression of the cell surface molecules CD58, CD80, CD83, CD86, HLA-ABC and HLA-DR (Fig. S7A and B) and the production of IL-12p70 (Fig. S7C). Thus, parasites alone had no effect on these maturation and activity markers of DCs but enhance innate immune responses. No such sensitization was observed with macrophages (Fig. S8).

### *Leishmania*-infected human DCs stimulate CD4+ and CD8+ T cells

Since human DCs handle *Leishmania* infection differently than macrophages, we investigated the capacity of *Leishmania*-infected DCs to induce CD4+ T helper and CD8+ cytolytic T cells. Day 5 iDCs were incubated for 48 h with either dead or live parasites and then in 7-days proliferation assays with autologous CFSE-labeled CD4+ and CD8+ T cells. Non-infected iDCs and DCs matured with LPS or cytokine cocktail (mDCs) were used as controls. As HLA restriction controls, we added pan–HLA class I-(W6/32) and class II–(TÜ39) specific mouse antibodies. Mouse anti-human Sec61 served as isotype control. All cells were from healthy donors in Germany without record of leishmaniasis. The T cells were thus naïve with respect to leishmanial antigens and the responses monitored in these experiments *in vitro* induced primary responses.

DCs treated with the proinflammatory cocktail alone were poor T cells stimulators, which is expected for autologous co-cultures of DCs and T cells. To further reduce potential non-leishmanial antigen-specific stimulation of T cells by autologous mDC, we used human instead of bovine serum in the cultures. DCs loaded with dead or infected with live parasites were poor stimulators of CD4+ T cells (Fig. S9 and Fig. 5A). On the other hand, DCs loaded with parasites plus a cocktail of inflammatory cytokines triggered vigorous CD4+ T cell proliferation (Fig. S9 and Fig. 5A). Efficient presentation of leishmanial antigens in the context of HLA class II appeared to depend on the maturation state of the DCs as neither dead nor live parasites alone were able to induce significant CD4+ T cell proliferation. In contrast, DCs infected with live parasites without inflammatory cytokines were efficient stimulators of CD8+ T cell proliferation, whereas DCs with dead parasites were good stimulators only when inflammatory cytokines were added (Fig. S9 and Fig. 5A). Likewise, production of IFN-γ was higher in co-cultures with DCs infected with live parasites (Fig. 5B). Inhibition of T cell proliferation and IFN-γ production with anti–HLA class I or II antibodies proved HLA restriction of the responses and rules out HLA-independent effects.

**Figure 5.**
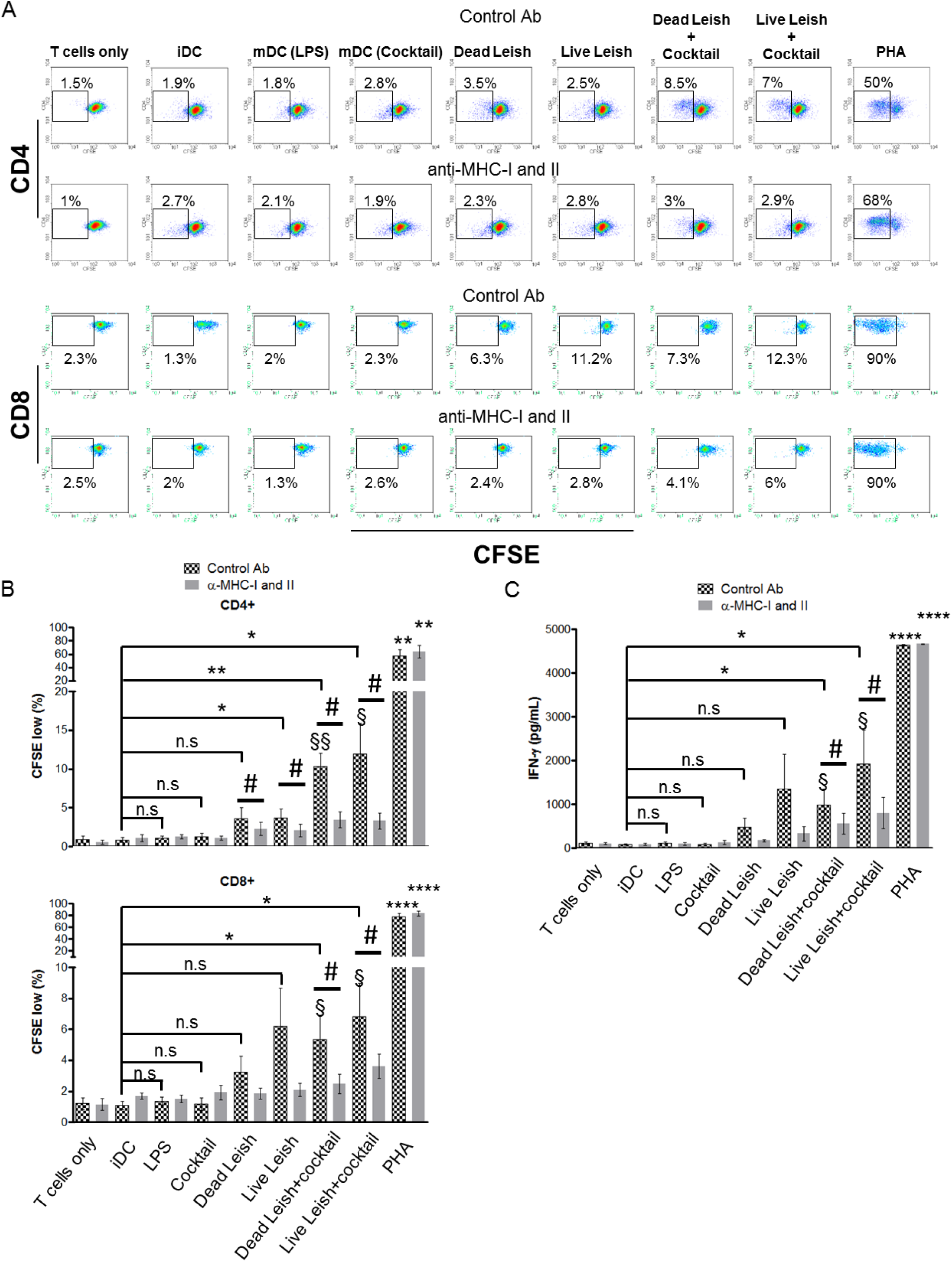
Induction of T cell responses by DCs infected with *L. donovani*. Adherent PBMCs were cultured in presence of GM-CSF and IL-4 for five days to generate iDCs that were matured with LPS or a cocktail of inflammatory cytokines (IL-1β, IL-6 and TNF-α), or cultured with dead or infected with live *L. donovani* parasites alone or together with the cocktail of inflammatory cytokines. After two days, the DCs were washed and characterized by flow cytometry as for the experiments shown in Figure S7 and then co-cultured with autologous CFSE-labeled CD3+ T cells. After seven days of co-culture, the cells were harvested and stained with antibodies against CD4 and CD8 and analyzed by flow cytometry. Proliferation of the T cells was determined by the reduction of CFSE upon successive cell divisions. Graphs in (**A**) show means of the percentage of CFSE^low^ cells among CD4 (upper graph) or CD8 (lower graph) from the five donors analyzed. The plots in (Fig S9) show CFSE fluorescence in CD4+ (upper panels) or CD8+ (lower panels) T cells of one of five independent experiments with cells from five different healthy donors. The supernatants of the co-cultures were collected and assayed for IFN-γ by ELISA (**B**). Cocktail: IL-1β, IL-6 and TNF-α. PHA: phytohaemagglutinin. n = 5. * = p<0.05; ** = p<0.01; and *** = p<0.001; **** = p<0.0001 comparing the groups treated as indicated with the control group iDC plus T cells (iDC) without parasites or cytokines or LPS. The T cell mitogen PHA served as positive control. # = p<0.05 comparing the group treated with control Ab vs the group treated with α-MHC-I and II.

## Discussion

Since immunity against leishmaniasis depends on CD4+ and CD8+ T cells rather than neutralizing antibodies ^28, 29^, DCs as key inducers of T cell responses are in the focus of the immunology of leishmaniasis and the development of anti-*Leishmania* vaccines and immunotherapies ^9,30–34^. Issues here involve up-take of the parasites, processing and presentation of their antigens, and induction of the T cells. That DCs efficiently take up *Leishmania* parasites like other phagocytes such as macrophages and neutrophils is expected and has been shown before^12^. However, that they, in contrast to the other phagocytes, destroy the parasites came as a surprise. It would have been expected that, if at all, macrophages as more aggressive phagocytes would be more efficient in that. But macrophages are the major host cells where *Leishmania* parasites proliferate until they rupture the cell and infect new macrophages. *Leishmania* parasites can be destroyed by CD8+ cytotoxic T cells inside infected macrophages via granulysin and granzyme B from their cytolytic granules, but this also kills the host cells ^35^. Moreover, the effector cells have to be induced first, which requires prior processing and presentation of leishmanial antigens by antigen-presenting cells, most importantly DCs^7^. Our data suggest that the destruction of intracellular parasites by their host DCs does not require the intervention by cytolytic T cells and is likely the link that leads to activation of effective *Leishmania*-specific adaptive and innate immune responses. The granulysin/granzyme B-dependent process is thought to be essential for the clearance of the parasites in human leishmaniasis, for cure from disease and for immunity to subsequent infections ^7, 35–37^. To our knowledge, this is the first report to show a microbicidal function of granulysin/granzyme B inside infected host DCs. It is worth noting that this mechanism cannot play a role in mice as mice do not have granulysin^8^, which makes it difficult to draw conclusions from murine models of leishmaniasis.

The destruction of the parasites in the DCs appears ultrastructurally like an apoptotic process, as the parasites undergo fragmentation. However, the molecular mechanisms seem to differ from the apoptosis of target cells induced by cytolytic T cells and involve, among other factors, reactive oxygen species triggered by the disruption of the respiratory chain ^23^. In effect, the destruction of the parasites inside the phagolysosomes would make their antigens accessible for the antigen processing machinery of the, and presentation by HLA class II and cross-presentation by HLA class I. This relates to the finding that the subcellular compartment that contains the parasites expresses MHC class I and class II molecules, as well as calnexin and invariant chain, chaperons that are required for the loading of the respective MHC molecules with peptides. ER-associated markers have been reported for phagosomes of murine DCs bearing *Toxoplasma gondii*^38^ where the ER-associated proteins were induced by live but not dead parasites. In our study, however, both dead and live parasites were found in compartments expressing calnexin. Other reports have shown recruitment of ER-derived vesicles by phagosomes of DCs that have taken in large particles^39^.

Since the destruction of the parasites by DCs follows an initial intracellular proliferation, the parasitocidal capacity seems to be induced by live but not dead parasites that may be processed like other particulate antigens. While this implies some degree of activation of the DCs, this does not extend to the well-established indicators of DC activation, CD83, the co-stimulatory molecules CD40, CD80 and CD86, and the cytokine IL-12. This again is surprising as components of *Leishmania* parasites have been reported to address innate pathogen-associated molecular pattern (PAMP) recognition receptors (PRRs) such as Toll-like receptors (TLRs), which should result in induction or up-regulation of said activation markers^40^. Maybe immunostimulatory leishmanial factors do not get access to the respective PRRs or these receptors are not expressed by iDCs. However, in the presence of proinflammatory cytokines, activation of *Leishmania*-bearing DCs is enhanced beyond what can be observed with the cytokines alone, which extends recently published observations with TLR agonists^41,42^. Some reports indicated that *Leishmania* alter DC maturation but the data are controversial, as some studies show induction of maturation by some species of *Leishmania* while others show impairment of DC maturation^12,40–43^. However, in the experiments that showed impairment of DC function, far higher MOIs than in our experiments were used. Obviously, when comparing different studies on the immunology of leishmaniasis careful attention needs to be paid to the detailed design of the different studies. In particular, it needs to be acknowledged that there are significant differences between different *Leishmania* species and the immune constitution of different host species.

Activation and maturation of DCs with expression of co-stimulatory molecules and IL-12 is essential for induction of efficient and sustained T cell responses. Several reports suggest that *Leishmania* parasites may suppress T cell induction. For instance, immature DC infected with *L. amazonensis* were shown to cause reduced production of IFN-γ by autologous T cells^44^. The authors of that report concluded that the parasites alter DC differentiation from human monocytes by down-regulating Th1-adaptative immune responses and, thereby, favoring their survival in the host cells. Our results contradict this conclusion. DCs incubated with dead or live parasites induce proliferation of CD4+ and CD8+ T cells, albeit weak, which, in case of the CD4+ T cells is strongly enhanced by proinflammatory cytokines that efficiently activate and drive maturation of the iDCs. In the case of CD8+ T cells, parasite-bearing DCs alone without additional cytokines induce strong CD8+ T cell proliferation, especially with DCs infected with live parasites. This latter response is as high as the responses seen in the presence of the cytokines. The T cells also produce IFN-γ in response to *Leishmania*-pulsed DC, which is well above background in the absence and enhanced in the presence of inflammatory cytokines. IFN-γ has important functions in anti-*Leishmania* immune responses, for instance for harnessing macrophages and as differentiation factor driving the development of effector T cells^45,46^. The T cell responses seen without addition of inflammatory cytokines also need activation and maturation of the DC, which may be conjectured to be driven by cognate interaction with T cells during the 7 days’ time period of the assay. Cognate interactions of DCs and T cells are, besides PAMP and inflammatory conditions, strong inducers of DC activation and maturation in particular in pre-immune individuals. Such cognate interactions of expanded *Leishmania*-specific T cell populations with *Leishmania*-bearing DCs may contribute to the immunity of pre-exposed individuals against leishmaniasis that is evidenced by the well documented immunity following recovery from diseases, which is in contrast to the situation with other protozoal diseases such as malaria^47,48^.

The data presented herein document a previously unknown mechanism for the control of *Leishmania* parasites by human DCs through granzyme B and granulysin. Furthermore, they emphasize the need to address the DCs when developing anti-*Leishmania* vaccines or immunotherapies in order to induce efficient CD4+ helper and CD8+ effector T cell responses. They may explain why leishmanization, i.e. immunization with live parasites, is efficient whereas subunit vaccines are not. However, leishmanization induces immunity through deliberate infection with subsequent disease, which may come with severe adverse effects. Alternative strategies may consider TLR agonists or inflammatory cytokines for *in situ* vaccination and immunotherapy to activate parasite-bearing DCs for induction of parasitocidal CD8+ effector T cell.

## Materials and Methods

### Human dendritic cells and macrophages

Dendritic cells (DCs) and macrophages were generated from plastic-adherent peripheral blood mononuclear cells (PBMCs) of healthy donors as described earlier ^49,50^. Briefly, adherent PBMCs were cultured in 75 cm^2^ cell culture flasks in RPMI 1640 GlutaMax culture medium (Invitrogen, Carlsbad, CA, USA) supplemented with 10% fetal bovine serum (FBS; Biochrom AG, Germany) at 37°C in humidified atmosphere with 8% CO_2_. For macrophages, the cultures were supplemented with 50 ng/mL recombinant human GM-CSF (Genzyme, Cambridge, MA, USA) on day 0 and 4. DC cultures were supplemented with 50 ng/mL IL-4 (PromoCell GmbH, Germany) in addition to GM-CSF. Where indicated, DC cultures included 1% converted human serum (Pan Biotech, Germany) instead of FBS. Live *L. donovani* were added at a multiplicity of infection (MOI) of 10:1 and dead parasites at the same ratio on day 5. Maturation of the DC was induced two hours after addition of the parasites with either 1 μg/mL LPS (Sigma-Aldrich, Germany) or a combination of IL-1β (10 ng/mL), IL-6 (25 ng/mL) and TNF-α (10 ng/mL) (all Strathmann Biotech, Hamburg, Germany). The DCs and macrophages were harvested at the indicated time points and processed for laser scanning confocal microscopy (LSCM), cell viability assays, phenotyping by flow cytometry or co-cultures with autologous T cells. DC supernatants were collected after 48 h for measurement of IL-12p70 or IFN-γ by ELISA. The use of human blood cells for the study were reviewed and approved by the institutional Ethics Committee of the Charité – Universitätsmedizin Berlin, Ethikausschuss 1 at Campus Charité Mitte (EA1/148/08 and EA1/026/14). The blood cells were obtained from the Institute of Transfusion Medicine of the Charité with informed consent of the donors for their use in research. All procedures were carried out in accordance with approved guidelines and regulations.

### Leishmania parasites

Promastigotes of the wild-type and YFP-transgenic *L. donovani* parasites strain BHU5 (MHOM/IN/02/BHU5)^51^ were cultivated at 25°C in M199 culture medium (Gibco Invitrogen) supplemented with 20% heat-inactivated FBS (Biochrom AG, Germany). For the experiments, the parasites were harvested at end-log-phase, centrifuged at 2800 rpm, suspended in RPMI 1640 GlutaMax culture medium (Invitrogen, Carlsbad, CA, USA), counted in a Neubauer chamber after fixation with 4% paraformaldehyde, and adjusted to the required densities for coculture with DCs or macrophages. Where dead parasites were used, they were killed with 4% paraformaldehyde for 15 min followed by extensive washing with PBS.

### Laser Scanning Confocal Microscopy

Uptake and routing of parasites in DCs, and granzyme B expression in DCs and macrophages and activity in DCs were monitored by Laser Scanning Confocal Microscopy (LSCM). To investigate the routing of the parasites into compartments containing MHC class I and II, invariant chain (CD74) and calnexin, DCs were harvested at the indicated time points and fixed with 4% paraformaldehyde for 20 min, permeabilized with 0.2% Triton X-100 in PBS for 10 min, incubated for 45 min with PBS-BSA 2% to block non-specific binding, and stained for 1.5 h with a monoclonal mouse IgG2a antibody against human HLA-DR, DP, DQ (BD Bioscience, Chicago, IL, USA), and rabbit IgG against calnexin or CD74 (both Santa Cruz Biotechnology, Santa Cruz, CA, USA). Alexa Fluor 647-labeled goat IgG anti-mouse IgG or anti-rabbit IgG (both Invitrogen, Carlsbad, CA, USA) were used as secondary antibodies. For analysis of the routing into MHC class I-positive compartments, the cells were stained with Alexa Fluor 647-labeled anti-HLA-A, B, C (BioLegend, San Diego, CA, USA) for 1 h. Fluorochrome-labelled secondary antibodies were used as controls for background fluorescence. To mark acidic compartments, live parasites-loaded DCs were incubated for 40 min with 100 nM LysoTracker Red DND-99 (Invitrogen, Carlsbad, CA, USA) followed by fixation with 4% paraformaldehyde for 20 min. Cells were subsequently counterstained with 4’,6-Diamidino-2-Phenylindole Dihydrochloride (DAPI; Roche Diagnostics GmbH, Mannheim, Germany) for nuclei visualization. To investigate granzyme B expression in DCs and macrophages, the cells were co-cultured with wild type parasites for 48 h were harvested and surface stained with APC-labelled anti-human CD11b and anti-human CD11c (both, BD Bioscience, Chicago, IL, USA), respectively. After surface staining, cells were fixed with Fix/Perm buffer (Invitrogen, Carlsbad, CA, USA) for 30 min, permeabilized with 1x permeabilization buffer (Invitrogen, Carlsbad, CA, USA) for 30 min and stained for 45 min with FITC-conjugated anti-human granzyme B (BD Bioscience, Chicago, IL, USA). To investigate granzyme B activity, DCs were co-cultured with YFP-transfected parasites with or without granzyme B fluorogenic substrate (Biomol GmbH, Hamburg, Germany) for 48 h, after which they were harvested and surface stained with APC-conjugated anti-human CD11c (BD Bioscience, Chicago, IL, USA). To determine the co-localization of parasites and granzyme B, DCs infected with YFP-transfected parasites for 48 h were harvested, fixed with Fix/Perm buffer (Invitrogen, Carlsbad, CA, USA) for 30 min, permeabilized with 1x permeabilization buffer (Invitrogen, Carlsbad, CA, USA) for 30 min and stained for 45 min with Alex Flour 647-conjugated anti-human granzyme B (BD Bioscience, Chicago, IL, USA). All stained cells were mounted on low-fluorescence slides in Fluoromount-G (SouthernBiotech) and covered with coverslips. The slides were analyzed by laser scanning confocal microscopy (Leica TCS SP2), and the images processed with the Leica Confocal Software Version 2.5 Build 1227.

### Gene expression using BioGPs

The gene expression profiles of granzyme B and granulysin in major human tissues including white blood cells, macrophages and DCs were extracted from BioGPS (http://biogps.org/) entries of curated micro array data from GeneAtlas U133A, gcrma.

### Gene expression analysis by RT-PCR

Total RNA was extracted from iDCs, mDCs, macrophages and PHA activated PBMCs using Nucleospin RNA II purification kit (Macherey-Nagel, Duren, Germany) as per the manufacturer’s instructions. 500 ng of DNase-treated RNA was used to prepare cDNAs using superscript III Reverse Transcriptase (Invitrogen, CA, USA). RT-PCR was carried out with 1 µl of cDNA with granzyme B (CTTCTGCTGGCCTTCCTCC and GACTTGGCTCCAGAGAAGGT), granulysin (AGGGTGTGAAAGGCATCTCA and AAGGACTACACAGCTCACCC) and ß-actin (AGGAGAAGCTGTGCTACGTC and CTCGTCATACTCCTGCTTGC) sequence-specific forward and reverse primers. PCR conditions were initial denaturation at 95 °C for 4 min followed by 35 cycles of 40 sec at 95 °C, 1 min at 50 °C to 68 °C, 40 sec at 72 °C and a final extension step of 10 min at 72 °C. PCR products were analyzed by 1% agarose gel electrophoresis. β actin was used as a control for the PCR reactions, PCR with cDNA from PHA-activated PBMCs and PCR minus cDNA were used as a positive and negative controls, respectively

### Inhibition of granzyme B and granulysin

To silence granzyme B and granulysin, iDCs were harvested from the cultures, washed three times with PBS and suspended in Accell Delivery Media without siRNA or with 1 μM siRNA for GrzB, GNLY or GrzB plus GNLY. The treated cells were incubated at 37 °C with 5% CO_2_ for 72 hours, washed three times with PBS, suspended in RPMI 1640 GlutaMax culture medium with 10% FCS and used for infection with YFP-transfected *L. donovani* parasites. To block the activity of granzyme B, iDCs were incubated with 100 µM granzyme B inhibitor (Ac-IETD-CHO) (Biochemicals, Hopkinton, MA) for 20 min and then co-cultured with YFP-transfected parasites for 24 or 48h.

### Autologous DC-T cell co-cultures

T lymphocytes (CD3+ T cells) were isolated from PBMCs by negative selection depleting all non-T cells: Monocytes, B and NK cells were stained with FITC or PE-labeled antibody against CD14, CD19 and CD56 (BD Bioscience, Chicago, IL, USA or Miltenyi Biotec GmbH, Bergisch Gladbach, Germany), respectively, and adsorbed to magnetic beads via antibodies against FITC or PE (Miltenyi Biotec GmbH, Bergisch Gladbach, Germany). These untouched CD3+ T cells were labeled with 10 μM CFSE (Invitrogen, Carlsbad, CA, USA) and cultured with autologous DCs (2×10^5^ T cells : 2×10^4^ DCs) in U-bottom 96 well plates for 7 days. Where indicated, anti-MHC class I (clone W6/32) and anti-class II (clone TÜ39; both BD Bioscience, Chicago, IL, USA) antibodies were added at 10 μg/mL. After co-culture, supernatants were harvested and measurement for IFN-γ by ELISA, and the cells stained with anti-CD4 and anti-CD8 antibodies for flow cytometry.

### Flow Cytometry

Granzyme B and granulysin expression in DCs and macrophages cultured with or without wild type parasites for 48 h was determined by flow cytometry after staining with APC anti-human CD11b or anti-human CD11c (both, BD Bioscience, Chicago, IL, USA), respectively, fixing with Fix/Perm buffer (Invitrogen, Carlsbad, CA, USA) for 30 min, permeabilizing with 1x permeabilization buffer (Invitrogen, Carlsbad, CA, USA) for 30 min, and staining for 45 min with FITC anti human granzyme B (BD Bioscience, Chicago, IL, USA) or Alexa fluor 647 anti granulysin (BioLegend, San Diego, CA, USA). For phenotype analyses, DCs were stained with fluorochrome-labeled monoclonal antibodies against CD1a, CD40, CD58, CD80, CD83, CD86, HLA-ABC and HLA-DR (all BD Bioscience, Heidelberg, Germany) and T lymphocytes with antibodies against CD4 and CD8 (BioLegend, San Diego, CA, USA or BD Bioscience, Heidelberg, Germany). Data were acquired using FACSCalibur flow cytometer (Becton Dickinson, Heidelberg, Germany), processed with the CellQuest software (Becton Dickinson, Heidelberg, Germany) and analyzed with the WinMDi 2.9 (Purdue University, USA) or FLOWJO, LLC (385 Williamson Way, Ashland, OR 97520, USA).

### Cytokine assays

IL-12p70 was measured in supernatants of DC cultures using a commercial ELISA kit (Max Standard kit; BioLegend, San Diego, CA, USA). IFN-γ in DC-T cell co-cultures was detected with a capture antibody against IFN-γ (Thermo Scientific, Schwerte, Germany; 5 μg/mL) in PBS (Gibco, Darmstadt, Germany) absorbed overnight at room temperature in 96-well microtiter Nunc Maxisorp plates (Rochester, NY, USA) and washed. After blocking with 1% BSA (Sigma-Aldrich, Steinheim, Germany) in PBS for 1 h at room temperature, samples (100 μL) were added and incubated for 90 min at room temperature. After washing, biotinylated detection antibody (2.5 μg/mL 1% BSA/PBS) was added for 60 min at room temperature. The plates were washed and streptavidin-conjugated alkaline phosphatase (R&D Systems, Wiesbaden, Germany) was added at a 1000-fold dilution, incubated for 30 min at room temperature and washed. pNPP (Sigma Diagostic, Steinheim, Germany) at 1 mg/ml 1 M Tris-Cl, pH 9.5 was added, and the plates were incubated for another 30 min at room temperature in the dark. Optical density at 405 nm was determined with an ELISA reader (Dynatech Laboratories, Denkendorf, Germany). All assays were performed in duplicates and the results are expressed as pg/mL.

### Cell viability assay

Cell viability was assessed by staining the cells with Calcein-AM (Invitrogen, Carlsbad, CA, USA) and propidium Iodide (PI) (Sigma-Aldrich, Germany). DCs or macrophages infected with *Leishmania* parasites were harvested at the indicated time points and labeled with calcein-AM, PI and antibodies anti-CD11c or anti Cd11b (both BD Bioscience, Chicago, IL, USA) for 20 min and analyzed by flow cytometry. The cells positive for calcein and negative for PI were considered alive and those positive for PI and/or negative for calcein dead.

### Statistical analysis

Data were first subjected to Kolmogorov-Smirnov normality tests and, when normally distributed, analyzed by the parametric t test. Data sets that did not pass the normality test were analyzed by the nonparametric Wilcoxon test. Results were expressed as the mean ± standard error (SEM). P < 0.05 was considered significant. Data were analyzed using GraphPad Prism 2.01 (San Diego, CA, USA).

## Author contributions

IMA, RBB and LWN performed the experiments and wrote the first draft of the manuscript. AK assisted in the gene expression analyses. PW proposed the project reported herein, supervised the work and wrote the manuscript together with IMA, RBB, LWN and AK.

## Acknowledgments

We gratefully acknowledge Shyam Sundar, Banares Hindu University, Varanasi, UP, India for the BHU5 parasites and Melanie Selka for providing the YFP-transgenic BHU5 parasites.

## Financial support

The work has been supported in parts by the BMBF-funded project IND 11/050 (Project number 01DQ12074), GHIT (Project number G2016-203) and institutional funds of the Charité.

## Potential conflict of interest

The authors declare no conflict of interest.

## Notes

### Competing Interest Statement

The authors have declared no competing interest.

## References

1. Ashford, R.W. Leishmaniasis reservoirs and their significance in control. Clin Dermatol 14, 523–32 (1996).

2. Alvar, J., Vélez, I.D., Bern, C., Herrero, M., Desjeux, P., Cano, J., Jannin, J., den Boer, M. WHO Leishmaniasis Control Team, Leishmaniasis worldwide and global estimates of its incidence. PLOS One 7, e35671 (2012).

3. Herwaldt, B.L. Leishmaniasis. Lancet 354, 1191–9 (1999).

4. Ho, M., Koech, D.K., Iha, D.W., Bryceson, A.D. Immunosuppression in Kenyan visceral leishmaniasis. Clin. Exp. Immunol. 51, 207–14 (1983).

5. Kevric, I., Cappel, M.A., Keeling, J.H. New World and Old World Leishmania Infections: A Practical Review. Dermatol. Clin. 33, 579–93 (2015).

6. Forestier, C.L., Gao, Q., Boons, G.J. Leishmania lipophosphoglycan: how to establish structure-activity relationships for this highly complex and multifunctional glycoconjugate? Front. Cell. Infect. Microbiol. 4, 193 (2015).

7. Stenger, S., Hanson, D.A., Teitelbaum, R., Dewan, P., Niazi, K.R., Froelich, C.J., Ganz, T., Thoma-Uszynski, S., Melián, A., Bogdan, C., Porcelli, S.A., Bloom, B.R., Krensky, A.M. An antimicrobial activity of cytolytic T cells mediated by granulysin. Science 282, 121–5 (1998).

8. Huang, L.P., Lyu, S.C., Clayberger, C., Krensky, A.M. Granulysin-mediated tumor rejection in transgenic mice. J. Immunol. 178, 77–84 (2007).

9. Liu, D., Uzonna, J.E. The early interaction of Leishmania with macrophages and dendritic cells and its influence on the host immune response. Front. Cell. Infect. Microbiol. 2, 83 (2012).

10. Steinman, R.M. Dendritic cells: understanding immunogenicity. Eur. J. Immunol. 37 (Suppl 1):S53–S60 (2007).

11. Banchereau, J., Steinman, R.M. Dendritic cells and the control of immunity. Nature 392, 245–52 (1998).

12. Ghosh, M., Mandal, L., Maitra, S., Rakshit, S., Paul, K., Bagchi, J., Ganguly, D., Pal, C., Bandyopadadhyay, S. Leishmania donovani infection of human myeloid dendritic cells leads to a Th1 response in CD4+ T cells from healthy donors and patients with kala-azar. J. Infect. Dis. 194, 294–301 (2006).

13. Ruiz, J.H., Becker, I. CD8 cytotoxic T cells in cutaneous leishmaniasis. Parasite Immunol. 29, 671–8 (2007).

14. van den Bogaart, E., de Bes, H.M., Balraadjsing, P.P., Mens, P.F., Adams, E.R., Grobusch, M.P., van Die, I., Schallig, H.D. Leishmania donovani infection drives the priming of human monocyte-derived dendritic cells during Plasmodium falciparum co-infections. Parasite Immunol. 37, 453–69 (2015).

15. Brodsky, F.M., Guagliardi, L.E. The cell biology of antigen processing and presentation. Annu. Rev. Immunol. 9, 707–744 (1991).

16. Neefjes, J., Ploegh, H.L. Intracellular transport of MHC class II Molecules. Immunol Today 13, 179–184 (1992).

17. Pasternack MS, Eisen HN. A novel serine esterase expressed by cytotoxic T lymphocytes. Nature. 1985 Apr 25-May 1;314(6013):743–5. doi: 10.1038/314743a0. PMID: 3873011.

18. Pasternack MS, Verret CR, Liu MA, Eisen HN. Serine esterase in cytolytic T lymphocytes. Nature. 1986 Aug 21-27;322(6081):740–3. doi: 10.1038/322740a0. PMID: 3489187.

19. Peña SV, Hanson DA, Carr BA, Goralski TJ, Krensky AM. Processing, subcellular localization, and function of 519 (granulysin), a human late T cell activation molecule with homology to small, lytic, granule proteins. J Immunol. 1997 Mar 15;158(6):2680–8. PMID: 9058801.

20. Peña SV, Krensky AM. Granulysin, a new human cytolytic granule-associated protein with possible involvement in cell-mediated cytotoxicity. Semin Immunol. 1997 Apr;9(2):117–25. doi: 10.1006/smim.1997.0061. PMID: 9194222.

21. Winkler U, Fraser SA, Hudig D. Perforin-enhancing protein, a low molecular weight protein of cytotoxic lymphocyte granules, enhances perforin lysis. Biochem Biophys Res Commun. 1997 Jul 9;236(1):34–9. doi: 10.1006/bbrc.1997.6899. PMID: 9223421.

22. Liu X, Lieberman J. Knocking ‘em Dead: Pore-Forming Proteins in Immune Defense. Annu Rev Immunol. 2020 Apr 26;38:455–485. doi: 10.1146/annurev-immunol-111319-023800. Epub 2020 Jan 31. PMID: 32004099; PMCID: PMC7260445.

23. Dotiwala F, Lieberman J. Granulysin: killer lymphocyte safeguard against microbes. Curr Opin Immunol. 2019 Oct;60:19–29. doi: 10.1016/j.coi.2019.04.013. Epub 2019 May 18. PMID: 31112765; PMCID: PMC6800608.

24. Junqueira C, Barbosa CRR, Costa PAC, Teixeira-Carvalho A, Castro G, Sen Santara S, Barbosa RP, Dotiwala F, Pereira DB, Antonelli LR, Lieberman J, Gazzinelli RT. Cytotoxic CD8+ T cells recognize and kill Plasmodium vivax-infected reticulocytes. Nat Med. 2018 Sep;24(9):1330–1336. doi: 10.1038/s41591-018-0117-4. Epub 2018 Jul 23. PMID: 30038217; PMCID: PMC6129205.

25. Walch M, Dotiwala F, Mulik S, Thiery J, Kirchhausen T, Clayberger C, Krensky AM, Martinvalet D, Lieberman J. Cytotoxic Cells Kill Intracellular Bacteria through Granulysin-Mediated Delivery of Granzymes. Cell. 2015 May 21;161(5):1229. doi: 10.1016/j.cell.2015.05.021. Epub 2015 May 21. PMID: 28910642.

26. Medzhitov, R. Toll-like receptors and innate immunity. Nat. Rev. Immunol. 1, 135–45 (2001).

27. Takeuchi, O., Akira, S. Pattern recognition receptors and inflammation. Cell 140, 805–20 (2010).

28. Kaye, P., Scott, P. Leishmaniasis: Complexity at the host–pathogen interface. Nat. Rev. Microbiol. 9, 604–615 (2011).

29. Maroof, A., Brown, N., Smith, B., Hodgkinson, M.R., Maxwell, A., Losch, F.O., Fritz, U., Walden, P., Lacey, C.N., Smith, D.F., Aebischer, T., Kaye, P.M. Therapeutic vaccination with recombinant adenovirus reduces splenic parasite burden in experimental visceral leishmaniasis. J. Infect. Dis. 205, 853–863 (2012).

30. Ashok, D., Schuster, S., Ronet, C., Rosa, M., Mack, V., Lavanchy, C., Marraco, S.F., Fasel, N., Murphy, K.M., Tacchini-Cottier, F., Acha-Orbea, H. Cross-presenting dendritic cells are required for control of Leishmania major infection. Eur. J. Immunol. 44, 1422–32 (2014).

31. Basu, R., Bhaumik, S., Haldar, A.K., Naskar, K., De, T., Dana, S.K., Walden, P., Roy, S. Hybrid cell vaccination resolves Leishmania donovani infection by eliciting a strong CD8+ cytotoxic T-lymphocyte response with concomitant suppression of interleukin-10 (IL-10) but not IL-4 or IL-13. Infect. Immun. 75, 5956–66 (2007).

32. Basu, R., Roy, S., Walden, P. HLA class I-restricted T cell epitopes of the kinetoplastid membrane protein-11 presented by Leishmania donovani-infected human macrophages. J. Infect. Dis. 195, 1373–80 (2007).

33. Das, S., Freier, A., Boussoffara, T., Das, S., Oswald, D., Losch, F.O., Selka, M., Sacerdoti-Sierra, N., Schönian, G., Wiesmüller, K.H., Seifert, K., Schroff, M., Juhls, C., Jaffe, C.L., Roy, S., Das, P., Louzir, H., Croft, S.L., Modabber, F., Walden, P. Modular multiantigen T cell epitope-enriched DNA vaccine against human leishmaniasis. Sci. Transl. Med. 6, 234ra56 (2014).

34. Freitas-Silva, R., Brelaz-de-Castro, M.C., Rezende, A.M., Pereira, V.R. Targeting Dendritic Cells as a Good Alternative to Combat Leishmania spp. Front. Immunol. 5, 604 (2014).

35. Dotiwala F, Mulik S, Polidoro RB, Ansara JA, Burleigh BA, Walch M, Gazzinelli RT, Lieberman J. Killer lymphocytes use granulysin, perforin and granzymes to kill intracellular parasites. Nat Med. 2016 Feb;22(2):210–6. doi: 10.1038/nm.4023. Epub 2016 Jan 11. PMID: 26752517; PMCID: PMC7325279.

36. Murray, H.W., Mitchell-Flack, M., Zheng, H., Ma, X. Granzyme-mediated regulation of host defense in the liver in experimental Leishmania donovani infection. Infect. Immun. 83, 702–12 (2015).

37. Stenger, S., Rosat, J.P., Bloom, B.R., Krensky, A.M., Modlin, R.L. Granulysin: a lethal weapon of cytolytic T cells. Immunol. Today 9, 390–4 (1999).

38. Goldszmid, R.S., Coppens, I., Lev, A., Caspar, P., Mellman, I., Sher, A. Host ER-parasitophorous vacuole interaction provides a route of entry for antigen cross-presentation in Toxoplasma gondii-infected dendritic cells. J. Exp. Med. 206, 399–410 (2009).

39. Cebrian, I., Visentin, G., Blanchard, N., Jouve, M., Bobard, A., Moita, C., Enninga, J., Moita, L.F., Amigorena, S., Savina, A. Sec22b regulates phagosomal maturation and antigen crosspresentation by dendritic cells. Cell 147, 1355–68 (2011).

40. Marovich, M.A, McDowell MA, Thomas EK, Nutman TB. IL-12p70 production by Leishmania major-harboring human dendritic cells is a CD40/CD40 ligand-dependentprocess. J. Immunol. 164, 5858–65 (2000).

41. Craft, N., Birnbaum, R., Quanquin, N., Erfe, M.C., Quant, C., Haskell, J., Bruhn, K.W. Topical resiquimod protects against visceral infection with Leishmania infantum chagasi in mice. Clin. Vaccine Immunol. 21, 1314–22 (2014).

42. Huang, L., Hinchman, M., Mendez, S. Coinjection with TLR2 agonist Pam3CSK4 reduces the pathology of leishmanization in mice. PLoS Negl. Trop. Dis. 9:e0003546 (2015).

43. Contreras, I., Estrada, J.A., Guak, H., Martel, C., Borjian, A., Ralph, B., Shio, M.T., Fournier, S., Krawczyk, C.M., Olivier, M. Impact of Leishmania Mexicana infection on dendritic cell signalling and functions. PLoS Negl. Trop. Dis. 8:e3202 (2014).

44. Favali, C., Tavares, N, Clarêncio, J, Barral, A, Barral-Netto, M, Brodskyn, C. Leishmania amazonensis infection impairs differentiation and function of human dendritic cell s. J. Leukoc. Biol. 82, 1401–6 (2007).

45. Lehmann, J., Enssle, K.H., Lehmann, I., Emmendörfer, A., Lohmann-Matthes, M.L. The capacity to produce IFN-gamma rather than the presence of interleukin-4 determines the resistance and the degree of susceptibility to Leishmania donovani infection in mice. J. Interferon Cytokine Res. 1, 63–77 (2000).

46. Nascimento, M.S., Carregaro, V., Lima-Júnior, D.S., Costa, D.L., Ryffel, B., Duthie, M.S., de Jesus, A., de Almeida, R.P., da Silva, J.S. Interleukin 17A acts synergistically with interferon γ to promote protection gainst Leishmania infantum infection. J. Infect. Dis. 211, 1015–26 (2015).

47. Abass, E., Kang, C., Martinkovic, F., Semião-Santos, S.J., Sundar, S., Walden, P., Piarroux, R., El Harith, A., Lohoff, M., Steinhoff, U. Heterogeneity of Leishmania donovani parasites complicates diagnosis of visceral leishmaniasis: comparison of different serological tests in three endemic regions. PLoS One 10(3):e0116408 (2015).

48. Das, S., Matlashewski, G., Bhunia, G.S., Kesari, S., Das, P. Asymptomatic Leishmania infections in northern India: a threat for the elimination programme? Trans. R. Soc. Trop. Med. Hyg. 108, 679–84 (2014).

49. Baleeiro, R.B., Rietscher, R., Diedrich, A., Czaplewska, J.A., Lehr, C.M., Scherließ, R., Hanefeld, A., Gottschaldt, M., Walden, P. Spatial separation of the processing and MHC class I loading compartments for cross-presentation of the tumor-associated antigen HER2/*neu* by human dendritic cells. OncoImmunology 4, 11:e1047585. eCollection (2015).

50. Baleeiro, R.B., Schweinlin, M., Rietscher, R., Diedrich, A., Czaplewska, J.A., Metzger, M., Lehr, C.M., Scherließ, R., Hanefeld, A., Gottschaldt, M., Walden, P. Nanoparticle-based mucosal vaccines targeting tumor-associated antigens to human dendritic cells. J. Biomed. Nanotech. 7, 1527–43 (2016).

51. Forgber, M., Basu, R., Roychoudhury, K., Theinert, S., Roy, S., Sundar, S., Walden, P. Mapping the antigenicity of the parasites in Leishmania donovani infection by proteome serology. PLoS One. 20, 1:e40 (2006).

