## Supplementary figures for "*Leishmania*-infected human dendritic cells clear their intracellular parasites by a mechanism involving the cytolytic effector molecules granzyme B and granulysin"

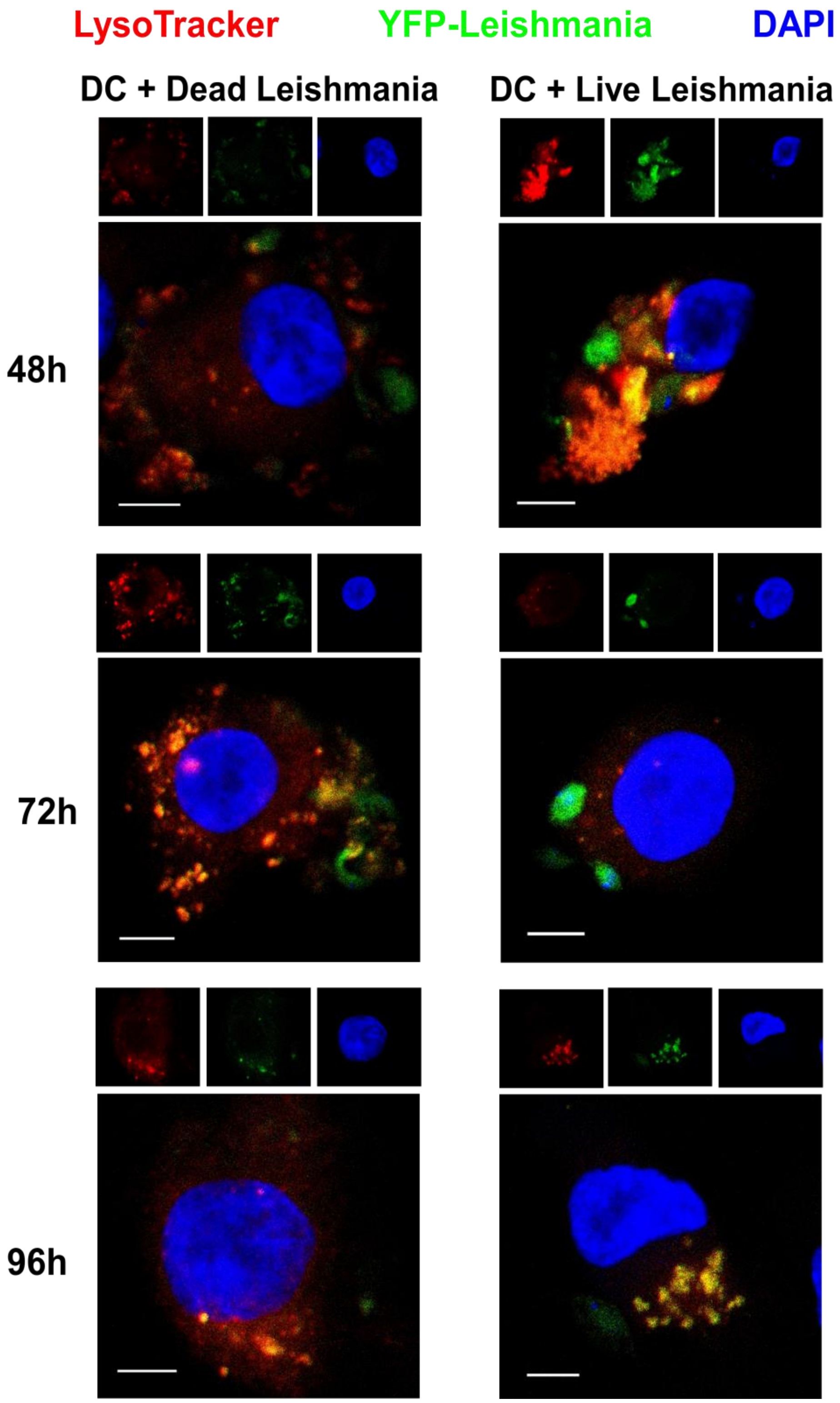


**Figure S1 | Kinetics of acidification of phagosomes containing *L. donovani* parasites.** Confocal images of day 5 iDCs incubated with YFP-transfected *L. donovani* parasites (green fluorescence) for the indicated period of time and stained with LysoTracker Red DND-99 (red fluorescence) for late/acidic endosomes, and DAPI (blue fluorescence) for nuclei. Parasites, or fragments thereof, inside acidic compartments appear in yellow. After staining, the cells were imaged by confocal microscopy. Scale bar= 5 μm.

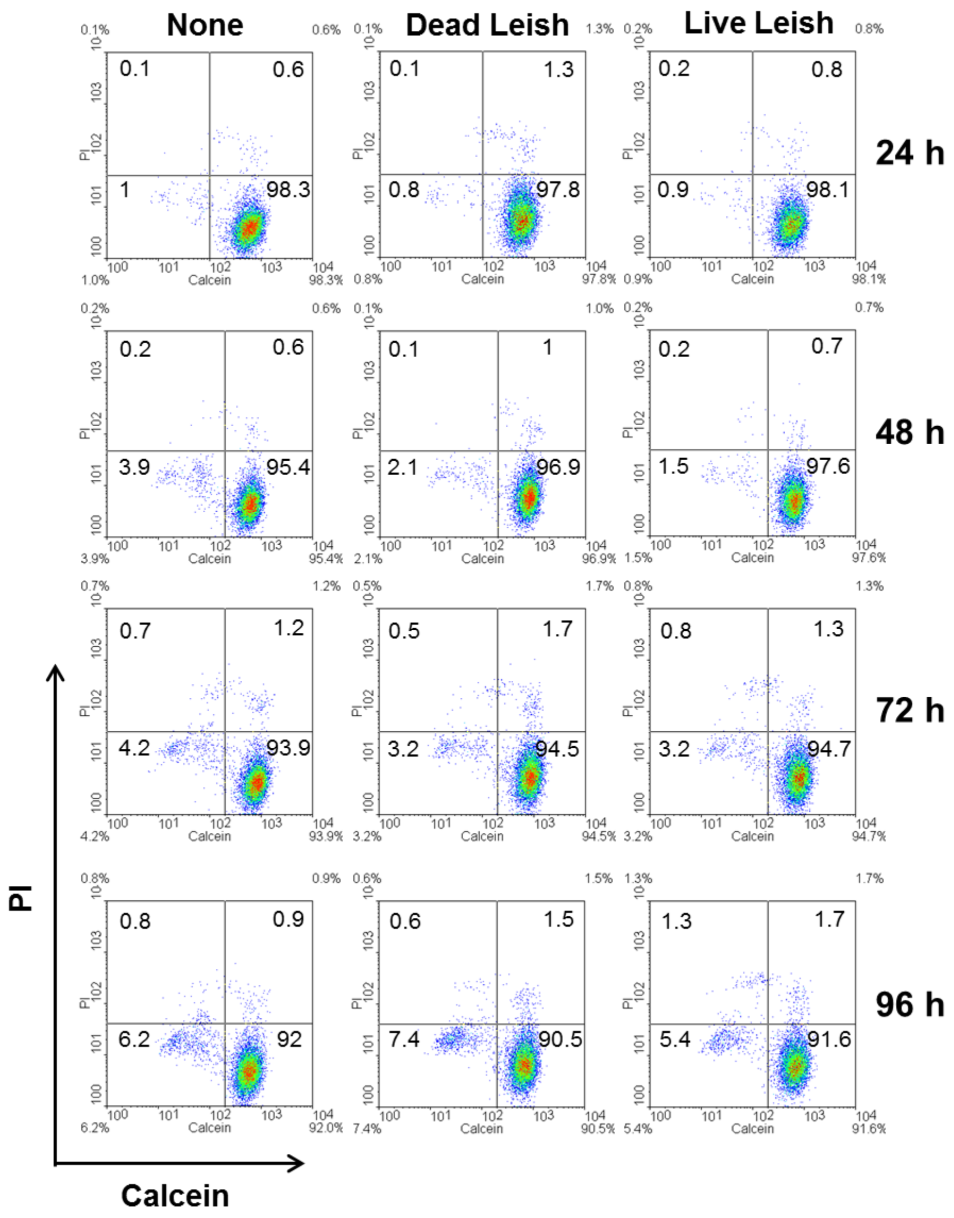


**Figure S2 | Effect of *L. donovani* on the viability of human iDCs.** The viability of the DCs was assessed after incubation with the parasites for the indicated period of time. DCs were stained with the nonfluorescent cell-permeable calcein-AM, which is metabolized in the cytoplasm of live cells into the green fluorescent calcein, and red fluorescent propidium iodide (PI), which only enters cells with perforated cell membranes and binds nucleic acids. After staining, the DC were analyzed by flow cytometry. Data shown are representative of four independent experiments performed with cells from four different donors.

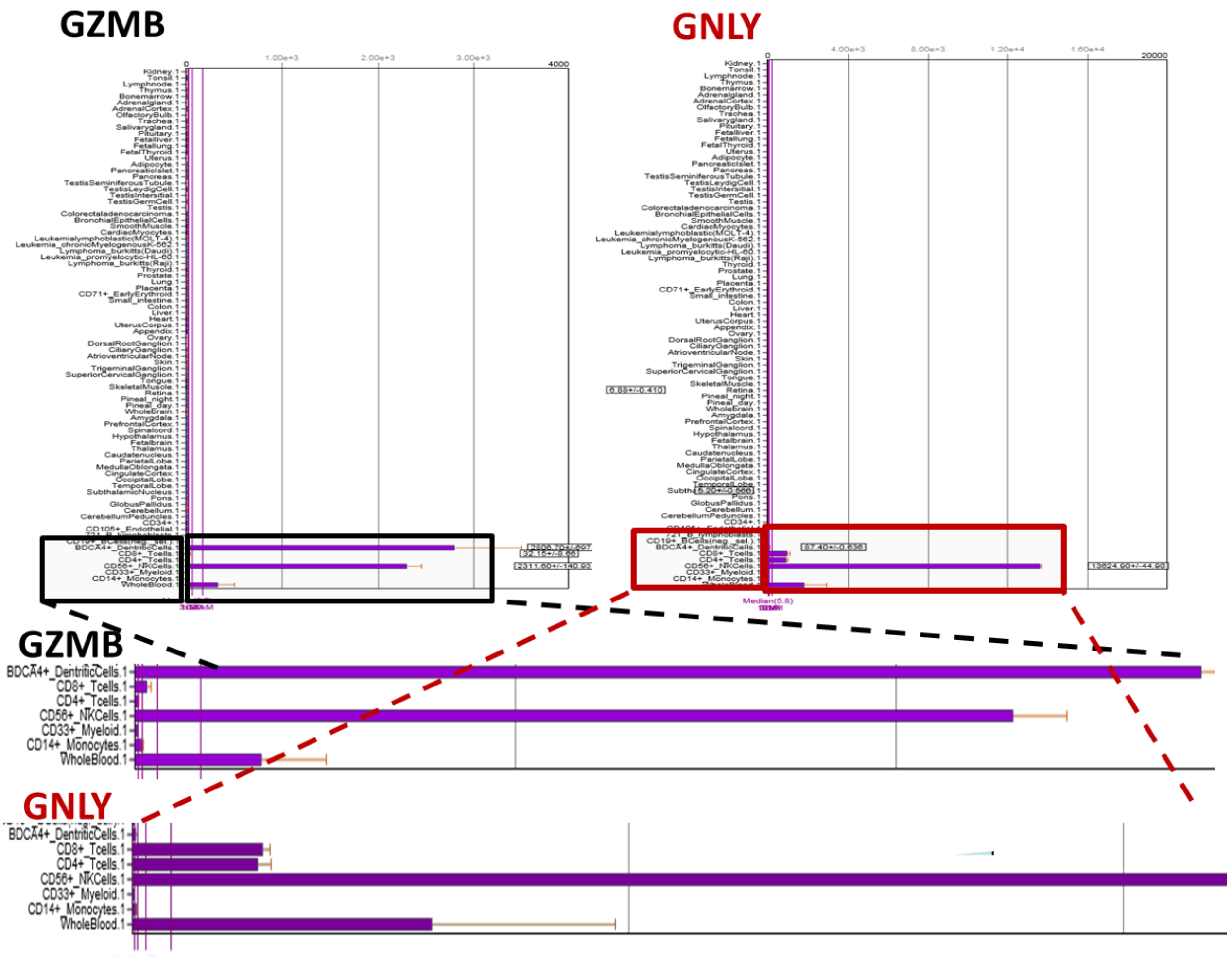


**Figure S3 | Granzyme B and granulysin gene expression using BioGPs.** The gene expression profiles of the granzyme B and granulysin in major human tissues, including human white blood cells. The BioGPS (http://biogps.org/) analyses were done with curated micro array data from GeneAtlas U133A, gcrma.

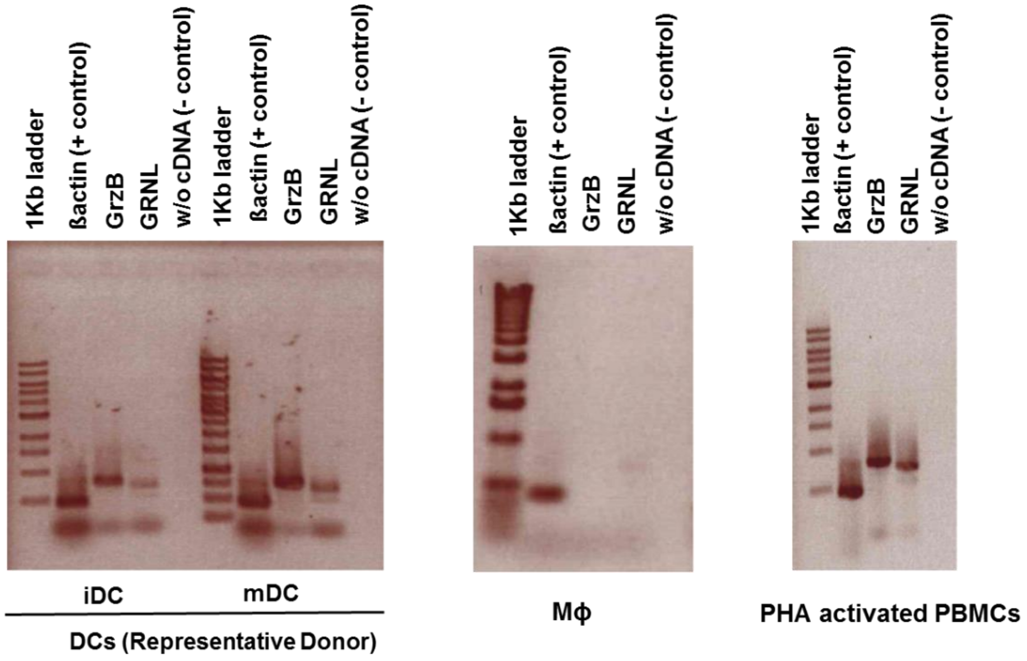


**Figure S4 | Expression of granzyme B and granulysin in iDCs, mDCs and macrophages (Mϕ)** RT-PCR showing expression of granzyme B and granulysin in iDCs, mDCs and macrophages. β actin was used as control for the PCR reactions. PCR with cDNA of PHA activated PBMCs and PCR minus cDNA were included as a positive and negative controls, respectively.

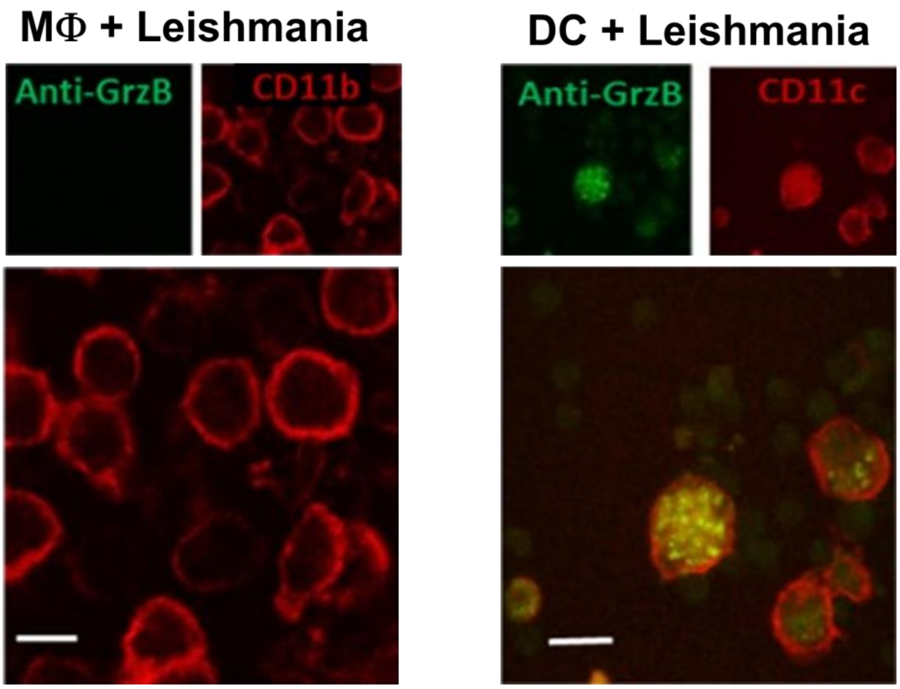


**Figure S5 | DC but not macrophages (MΦ) express granzyme B following infection with *L. donovani* parasites.** Day 8 MΦ and day 5 iDCs were infected with live *L. donovani* parasites for 48h, harvested and surfaced stained with fluorochrome-conjugated anti-CD11b or anti-CD11c, respectively. After cell-surface staining, cells were permeabilized and intracellularly stained with fluorochrome-conjugated anti-human granzyme B monoclonal antibodies analyzed by confocal microscopy. Image on the left depict macrophages with no detectable granzyme B while DCs shown in the image on the right exhibit granzyme B expression. CD11b and CD11c appear in red and identify macrophages and DC, respectively; and granzyme B appears in green. Scale bar= 5 µm.

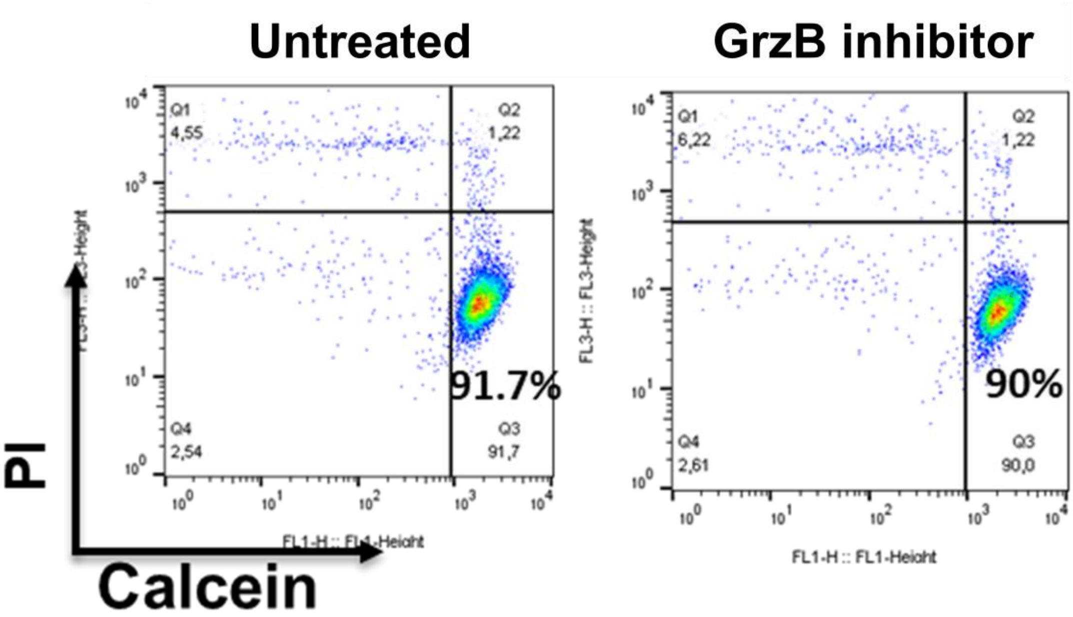


**Figure S6 | Treatment of DC with GrzB inhibitor does not affect their viability.** Day 5 iDCs were left untreated or were treated with GrzB inhibitor for 48h and their viability was analyzed by staining with calcein-AM and PI followed by flow cytometry.

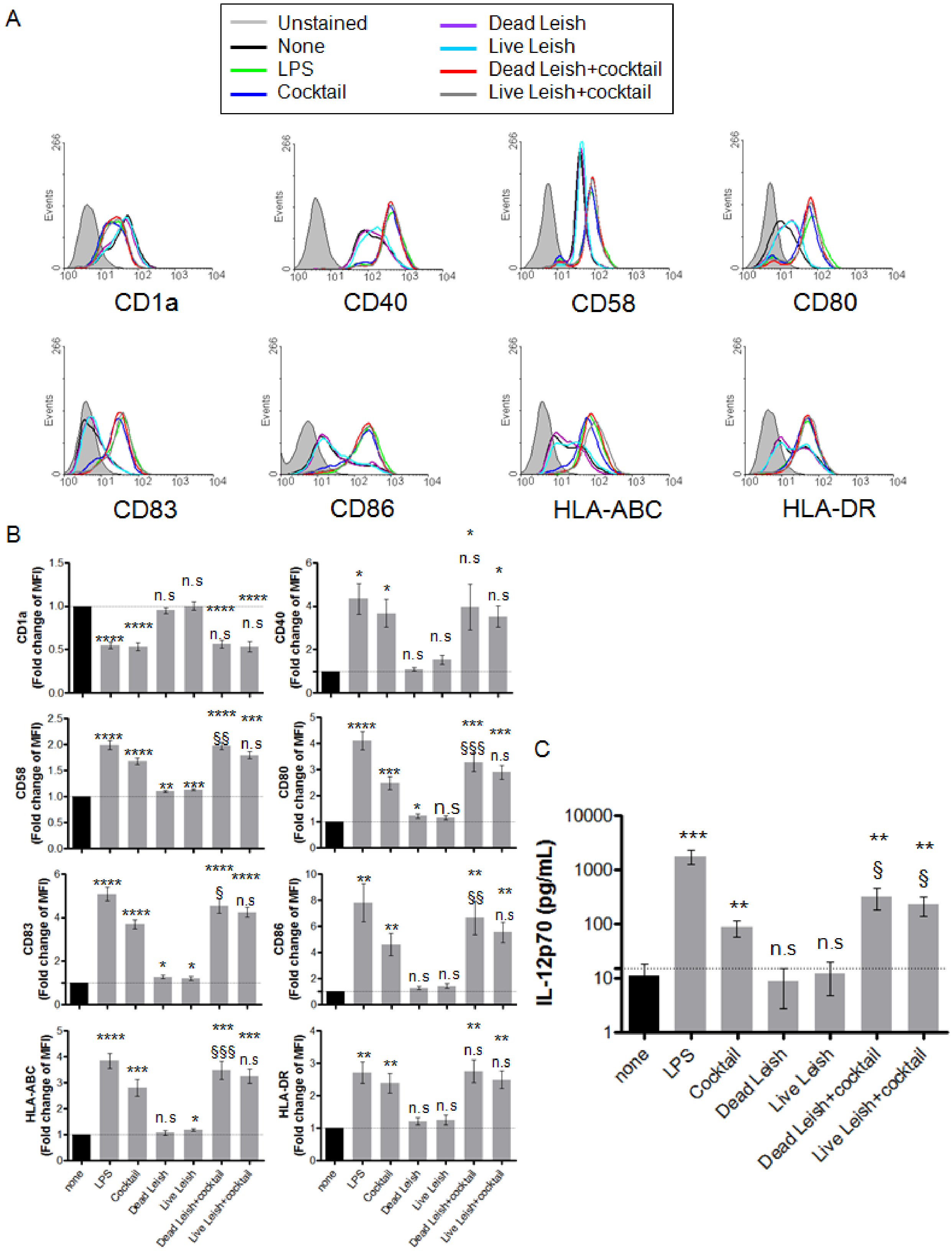


**Figure S7 | Effect of *L. donovani* infection on the activation and maturation states of DCs.** iDCs were generated from adherent human PBMCs cultured in RPMI 1640 medium supplemented with 10% FCS plus GM-CSF and IL-4. On day 5, the cells were harvested and incubated with LPS, an inflammatory cytokine cocktail of IL-1β, IL-6 and TNF-α, dead or live parasites, and dead or live parasites plus the cytokine cocktail for 2 additional days. On day 7, the cells were stained with fluorochrome-conjugated antibodies against CD1a, CD40, CD58, CD80, CD83, CD86, HLA-ABC and HLA-DR and analyzed by flow cytometry. The histograms show the expression of the markers by cells from one representative donor (**A**), and the graphs in (**B**) mean fold changes with standard error of the expression of the markers from the 8 donors analyzed. Supernatants were collected and assayed for IL-12p70 by ELISA (n=11) (**C**). Cocktail: IL-1β, IL-6 and TNF-α; n.s. = non-significant; * = p<0.05; ** = p<0.01; and *** = p<0.001; **** = p<0.0001 comparing the groups treated with the indicated factors vs. the control group "none". § = p<0.05; §§ = p<0.01; and §§§ = p<0.001 comparing the group treated with the indicated factors vs. the group "cocktail".

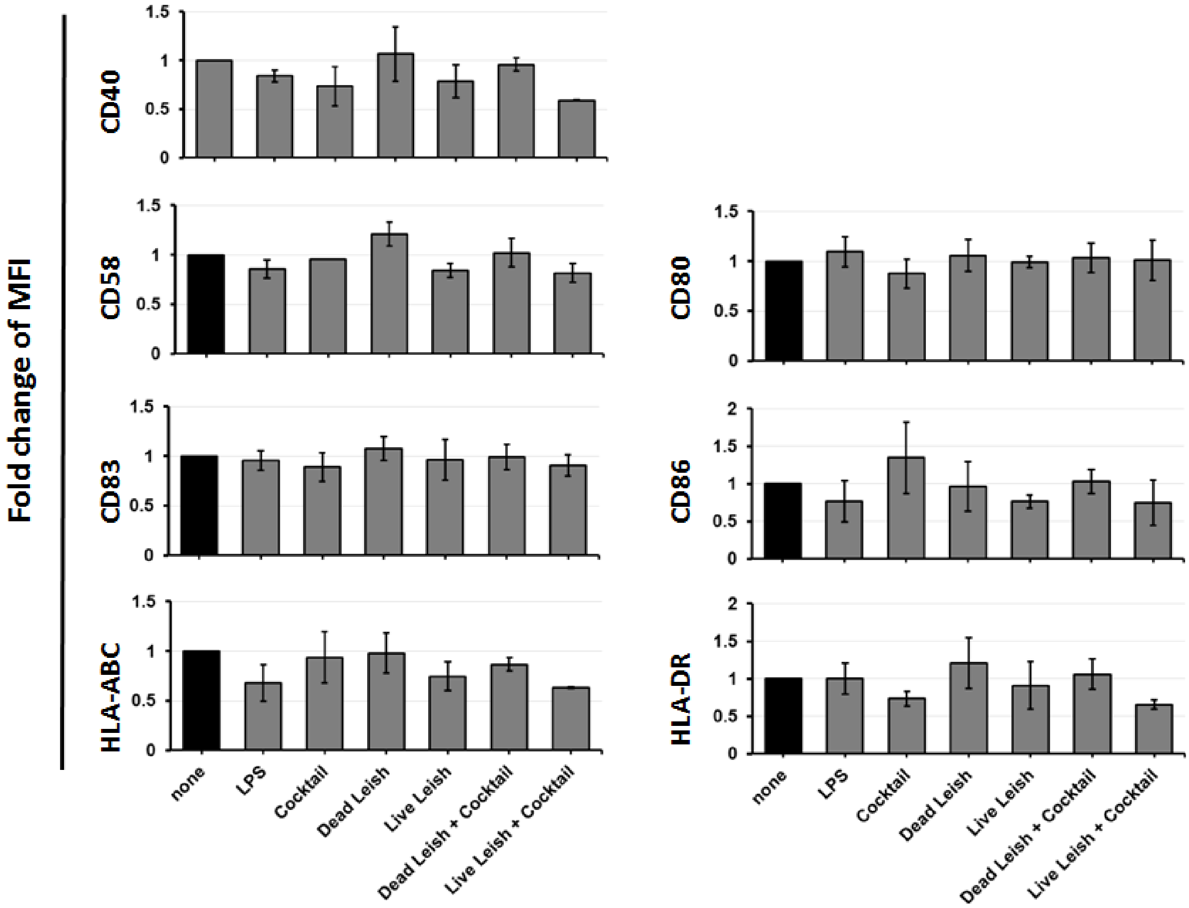


**Figure S8 | Effect of *L. donovani* infection on the activation of macrophages (MΦ).** Macrophages were generated from adherent PBMCs as described in material and methods. On day 8, the cells were harvested and incubated with LPS, an inflammatory cytokine cocktail of IL-1β, IL-6 and TNF-α, dead or live parasites, and dead or live parasites plus the cytokine cocktail for 2 additional days. On day 10 the cells were stained with fluorochrome-conjugated antibodies against CD40, CD58, CD80, CD83, CD86, HLA-ABC and HLA-DR and analyzed by flow cytometry. The graphs show mean fold changes with standard error of the expression of the markers from the 4 donors analyzed.

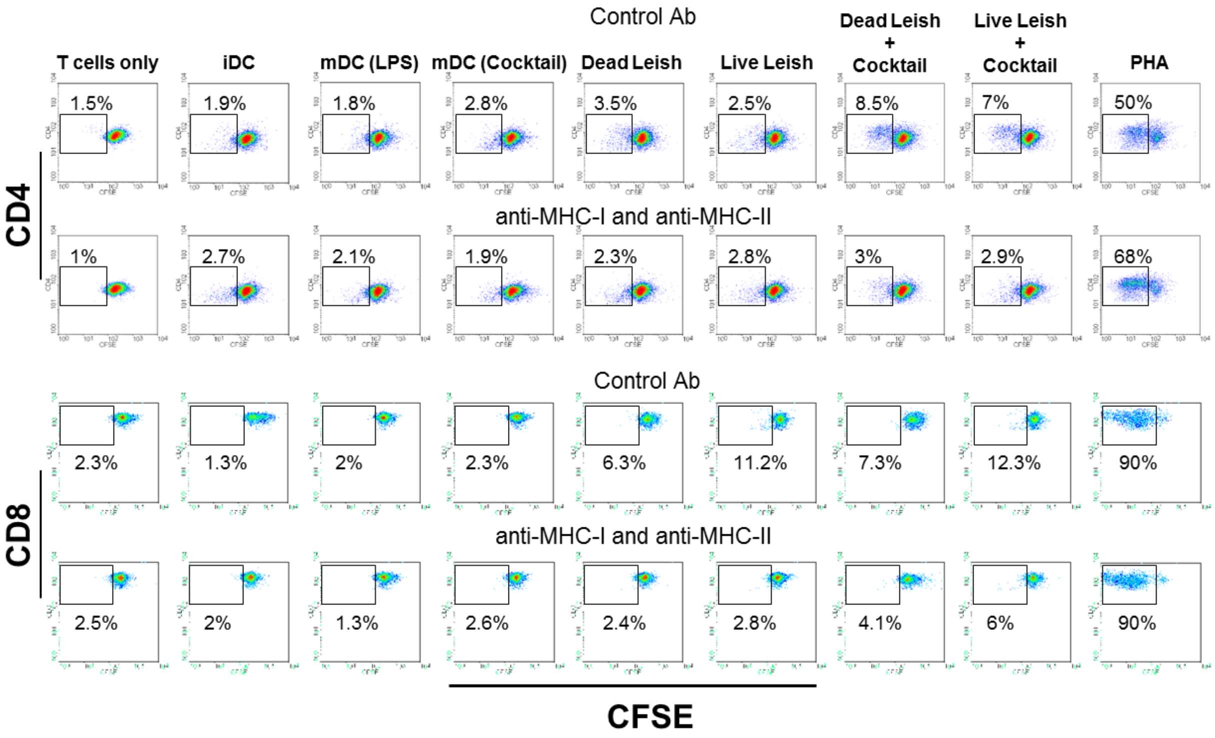


**Figure S9 | Induction of T cell responses by DC infected with *L. donovani*.** Adherent human PBMCs were cultured in presence of GM-CSF and IL-4 for five days and matured with LPS, a cocktail of inflammatory cytokines (IL-1β, IL-6 and TNF-α) or cultured with dead or infected with live *L. donovani* parasites alone or together with the cocktail of inflammatory cytokines. After two days, the DC were harvested, washed and characterized by flow cytometry and then co-cultured with autologous CFSE-labeled CD3+ T cells. After seven days of co-culture, the cells were harvested and stained with antibodies to CD4 and CD8 and analyzed by flow cytometry. Proliferation of the T cells was determined by the reduction of CFSE upon successive cell divisions. The plots show CFSE and CD4 (upper panels) or CD8 (lower panels) on T cells of one out of five independent experiments with cells from five different healthy donors.
